# NDH complex-mediated cyclic electron flow in bundle sheath cells enables C_4_ photosynthesis

**DOI:** 10.1101/2023.09.17.558135

**Authors:** Maria Ermakova, Russell Woodford, Duncan Fitzpatrick, Soraya M. Zwahlen, Graham Farquhar, Susanne von Caemmerer, Robert T. Furbank

## Abstract

The superior productivity of C_4_ plants is achieved via a metabolic C_4_ cycle which acts as a CO_2_ pump across mesophyll and bundle sheath (BS) cells and requires an additional input of energy in the form of ATP. Chloroplast NADH dehydrogenase-like complex (NDH) increases ATP production in C_3_ plants by operating cyclic electron flow (CEF) around Photosystem I (PSI), and its importance for C_4_ photosynthesis has been proposed from evolutionary and reverse genetics studies. We used the gene-edited C_4_ species *Setaria viridis* with null *ndhO* alleles lacking NDH to study a contribution of the complex to the cell-level electron transport. Our results indicate that NDH is the primary PSI electron acceptor mediating the majority of CEF in BS cells whilst the contribution of the complex to CEF in mesophyll cells is minimal. Moreover, the reduced leaf CO_2_ assimilation rate and growth of plants lacking the complex cannot be rescued by supplying additional CO_2_, indicating that NDH is essential for generating ATP required for CO_2_ fixation by the C_3_ cycle. Hereby we resolve a cell-level mechanism for the contribution of NDH to supporting high CO_2_ assimilation rates in C_4_ photosynthesis.

## Introduction

C_4_ plants typically exhibit superior radiation, water and nitrogen use efficiency allowing them to outperform C_3_ plants in warm climates^1,2^. The C_4_ cycle operates across mesophyll and bundle sheath (BS) cells as a biochemical carbon concentrating mechanism (Fig. 1). This increases CO_2_ partial pressure in BS cells where Rubisco and other Calvin-Benson-Bassham cycle (hereafter, the C_3_ cycle) enzymes reside^3^. The C_4_ cycle begins in the mesophyll cytosol with the conversion of CO_2_ to HCO_3_ ^-^ by carbonic anhydrase which is then fixed by PEP carboxylase (PEPC) into the C_4_ acid oxaloacetate. The latter can be reduced to malate or transaminated to aspartate before diffusing to BS cells for decarboxylation. There are three major subtypes of C_4_ photosynthesis, categorised by the primary decarboxylating enzyme^4^. As the majority of agriculturally important C_4_ species such as maize, sugarcane, sorghum and millets, use NADP^+^-dependent malic enzyme (NADP-ME) to decarboxylate malate in BS chloroplasts, this subtype is the major focus of C_4_ photosynthesis research and of this study. Decarboxylation of malate releases CO_2_, reduces NADP^+^ to NADPH and produces pyruvate which returns to mesophyll cells and is regenerated into PEP to complete the C_4_ cycle. Engineering C_3_ plants to carry out C_4_ photosynthesis is considered a promising route to increasing crop productivity, prompting attempts to introduce the C_4_ pathway into the C_3_ plant rice^5,6^. However, these projects are hampered by our incomplete understanding of the molecular mechanisms of C_4_ photosynthesis. Electron transport reactions crucial for providing at least two extra ATP molecules for each CO_2_ fixed – the additional energy cost of the carbon concentrating mechanism — is one such mechanism we aim to address in this work.

**Fig. 1.**
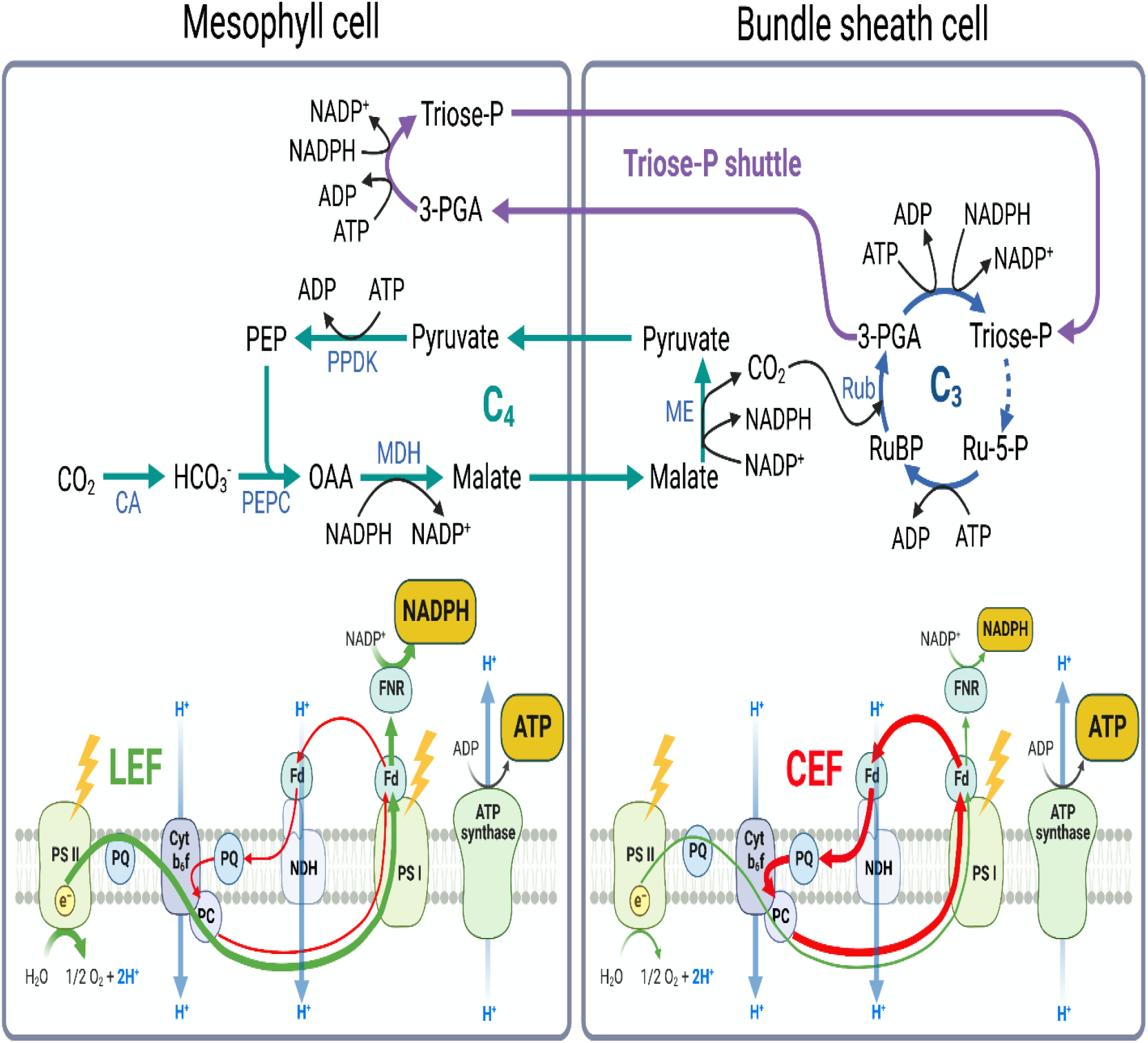
Schematic representation of metabolic (top part) and light (bottom part) reactions of C_4_ photosynthesis. CA, carbonic anhydrase; PEP, phosphoenolpyruvate; PEPC, phosphoenolpyruvate carboxylase; PPDK, pyruvate phosphate dikinase; OAA, oxaloacetate; MDH, malate dehydrogenase; ME, NADP^+^-dependent malic enzyme; RuBP, ribulose-5-bisphosphaste; Rub, Rubisco; 3-PGA, 3-phospho-glyceraldehyde; Triose-P, triose phosphate; Ru-5-P, ribulose-5-phosphate; LEF, linear electron flow; CEF, cyclic electron flow; PSII, Photosystem II; PQ, plastoquinone; Cytb_6_f, Cytochrome *b*_6_*f* complex; PC, plastocyanin; NDH, chloroplast NADH dehydrogenase-like complex; Fd, ferredoxin; PSI, Photosystem I; FNR, ferredoxin:NADP^+^ oxidoreductase.

In the C_4_ system, the electron transport chains of mesophyll and BS cells are tailored for specific metabolic needs^7^ (Fig. 1). The mesophyll electron transport chain closely resembles that of C_3_ plants, where linear electron flow (LEF) from Photosystem II (PSII) to Photosystem I (PSI) results in production of NADPH. Cytochrome *b*_6_*f* complex (Cyt*b*_6_*f*) between the two photosystems couples electron transport with proton translocation across the thylakoid membrane, which establishes a proton motive force (*pmf*) used by ATP synthase to generate ATP. For the C_4_ cycle, NADPH produced in mesophyll cells is primarily used for reducing oxaloacetate to malate while the ATP produced here is used to regenerate PEP; both are also used for the generation of triose phosphate within the C_3_ cycle (see below). The ΔpH component of *pmf* is also a key signal controlling electron transport rate by (i) slowing down Cyt*b*_6_*f* activity to restrict electron flow to PSI, also known as ‘photosynthetic control’^8^, and (ii) initiating a dissipation of absorbed light energy as heat through the rapidly-formed and reversible form of non-photochemical quenching (NPQ)^9^. This form of NPQ is activated by protonating lumen-exposed residues of the PsbS protein^10^ and through the conversion of violaxanthin to zeaxanthin in the antennae^11^.

LEF alone is often unable to satisfy the combined ATP/NADPH requirements of photosynthesis and other metabolic processes^12^. The C_3_ cycle in BS cells needs 3 mols of ATP and 2 mols of NADPH to fix 1 mol of CO_2_. While malate decarboxylation provides at least half of the required NADPH, mesophyll electron transport also supplies NADPH and ATP through the triose phosphate shuttle in which a part of the 3-PGA produced by Rubisco diffuses to the mesophyll cells for conversion to triose phosphate, which then returns to the BS^3,13^ (Fig. 1). This results in a lower requirement to produce NADPH in BS cells in which abundance and activity of PSII are diminished compared to the mesophylI^14^. It has long been proposed that to predominantly yield ATP, BS cells maintain active cyclic electron flow (CEF) that returns electrons from the reducing side of PSI back to the plastoquinone pool, allowing electrons to again pass through Cyt*b*_6_*f* and increase *pmf*^15,16^. There are two major proposed CEF pathways: via PROTON GRADIENT REGULATION5 (PGR5) and via the chloroplast NADH dehydrogenase-like complex (NDH)^17^. Abundance of both PGR5 and NDH in leaves increased during the evolutionary transition from C_3_ to C_4_ plants^18,19^ and NDH subunits preferentially accumulated in BS cells^20-22^. In C_3_ plants, NDH comprises only 0.2% of total thylakoid protein content^23^ and mutants lacking the complex retain normal fitness under optimal growth conditions^24^. In contrast, lowering NDH content in C_4_ species maize and *Flaveria bidentis* had severe effects on photosynthesis and growth^25-27^. While a direct mechanism for this effect has not been resolved, these observations prompted suggestions that NDH is involved in ATP production via CEF^26^ required for concentrating CO_2_ in BS cells^25^. Here we use CRISPR/Cas9 in a model C_4_ plant of NADP-ME subtype *Setaria viridis* to create plants lacking NDH and provide the first functional demonstration of NDH activity in C_4_ BS cells.

## Results

### Creating *S. viridis* plants with null *ndhO* alleles

To create null *ndhO* alleles, known to result in the arrested assembly of NDH^25^, we targeted Cas9 to the third and the fifth exons of *S. viridis ndhO* (Fig. 2a). Two new *ndhO* alleles were obtained: *ndhO-2* with a single nucleotide insertion causing a frameshift mutation, resulting in an altered amino acid (aa) sequence starting from S23 and a premature termination codon, and *ndhO-6* with a single nucleotide deletion causing a frameshift mutation, resulting in an altered aa sequence starting from G67 (Fig. S1). *S. viridis* plants with homozygous *ndhO-2* and *ndhO-6* alleles (*ndh* hereafter) lacked NDH as confirmed by immunoblotting of leaf extracts with antibodies against the NdhH subunit of the complex (Fig. 2b). In the absence of NDH, plants showed severe reduction of the aboveground biomass to about 30% of wild type (WT, Fig. 2d), which could not be rescued by supplementing air in the growth room with 2% CO_2_, a treatment previously shown to improve growth of *S. viridis* deficient in carbon concentrating mechanism^28,29^. The CO_2_ response of CO_2_ assimilation rate in *S. viridis ndh* plants closely resembled the maize *ndhO* mutant^25^, with assimilation reduced to about 50% of WT-level at intercellular CO_2_ partial pressures above 100 μbar (Fig. 2e). No differences between *ndhO-2* and *ndhO-6* plants indicated that both edited alleles resulted in inactive NDH.

**Fig. 2.**
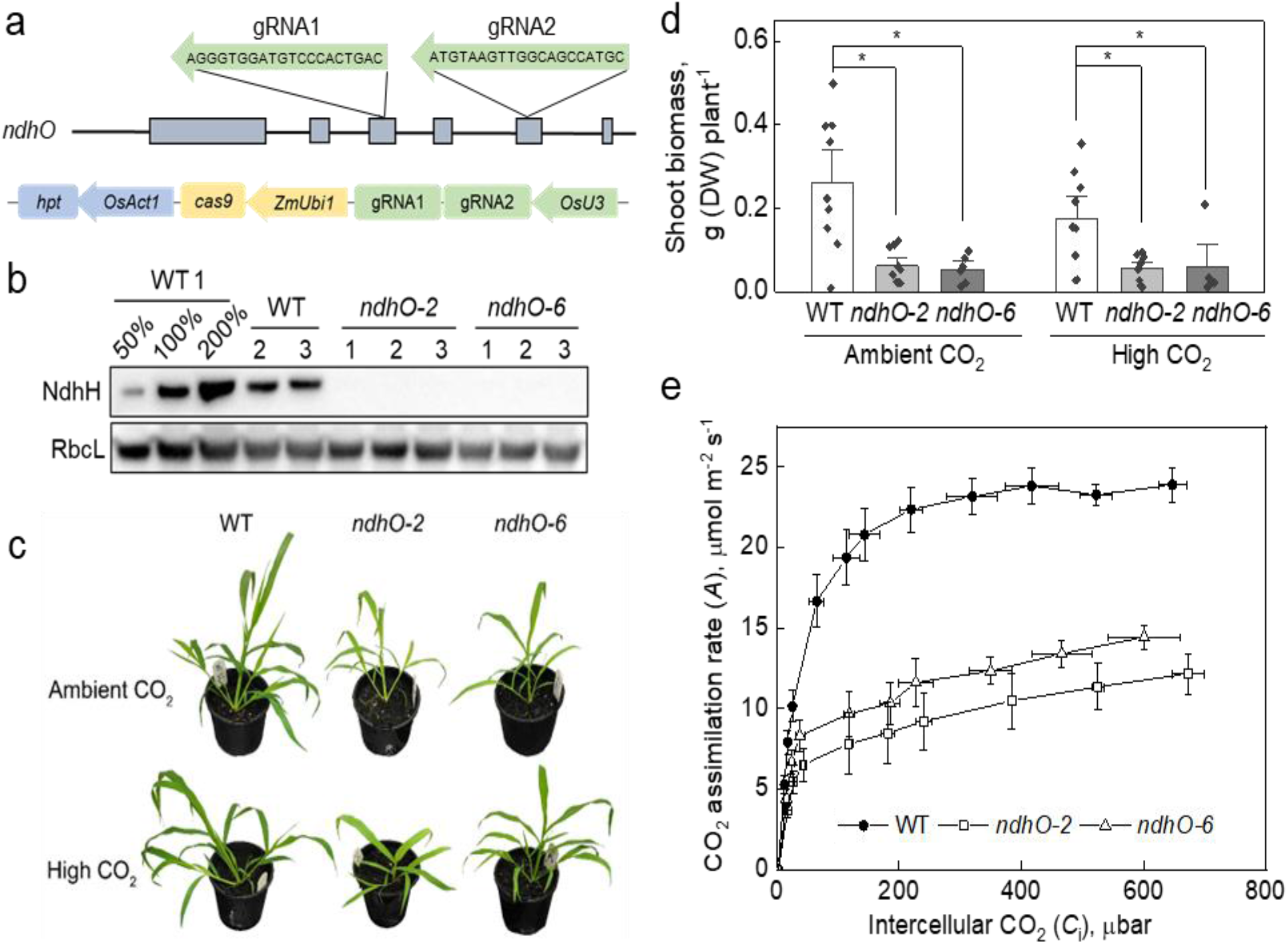
Creating *S. viridis* plants lacking NDH complex. **a**. Positions of guide RNAs (gRNAs) within the third and fifth exons of the *ndhO* genomic sequence and schematics of the gene construct assembled for transformation (see details in Materials and Methods). **b**. Immunodetection of the NdhH subunit of NDH and the large subunit of Rubisco (RbcL) in leaf protein extracts from wild-type (WT) and *ndhO-2* and *ndhO-6* plants with homozygous null *ndhO* alleles loaded on leaf area basis. Relative quantification of the blots is shown in Fig. S2. (**c, d**) Phenotype and biomass of plants grown for two weeks in air (Ambient CO_2_) or in air with 2% CO_2_ (High CO_2_). **d**. Mean ± SE, points are biological replicates. Asterisks indicate significant differences between edited and WT plants (*P* = 0.00003 for *ndhO-2, P* = 0.0002 for *ndhO-6*), no differences were found between *ndhO-2* and *ndhO-6* plants (*P* = 0.99) or between the CO_2_ treatments (*P* = 0.24) using two-way ANOVA and Tukey’s *post-hoc* test at α = 0.05. **e**. Response of CO_2_ assimilation rate (*A*) to intercellular CO_2_ partial pressures (*C*_i_) measured at 1500 μmol photons m^-2^ s^-1^ from plants grown at ambient CO_2_. Mean ± SE, *n* = 5 biological replicates. *A* was significantly decreased in both *ndhO-2* and *ndhO-6* plants compared to WT at *C*_i_ above 100 μbar (one-way ANOVA and Tukey’s *post-hoc* test at α = 0.05).

### Content of chlorophyll and photosynthetic proteins

Leaf contents of metabolic enzymes, PEPC, the large subunit of Rubisco (RbcL) and sedoheptulose-bisphosphatase (SBPase) of the C_3_ cycle, were unaltered in *ndh* plants (Fig. 2b, Fig, S2a). Relative abundances of some electron transport components, the D1 subunit of PSII, the AtpB subunit of ATP synthase and PsbS, did not differ between the gene-edited and WT plants per leaf area. However, plants lacking NDH had significantly less of the Rieske subunit of Cyt*b*_6_*f*, the PsaB subunit of PSI and PGR5 per leaf area, compared to WT (Fig. S2b). The total leaf Chl content was about 30% lower in *ndh* plants (Table S1). Mesophyll Chl content per leaf area was also significantly decreased in plants lacking NDH, whilst the BS Chl content was unaltered, compared to WT (Table S1).

The overall composition of thylakoid membranes was largely unaffected in mesophyll and BS cells of *ndh* plants, compared to WT (Fig. S2c), but cell-level changes in abundances of some electron transport components were detected by immunoblotting (Fig. 3). NDH complex was predominantly found in BS cells of WT plants. Relative contents of AtpB, PGR5 and PsbS per Chl were increased in BS cells of *ndh* plants (Fig. 3b) whilst Rieske and PGR5 abundance was significantly decreased in mesophyll cells of *ndhO* plants, compared to corresponding WT cells.

**Fig. 3.**
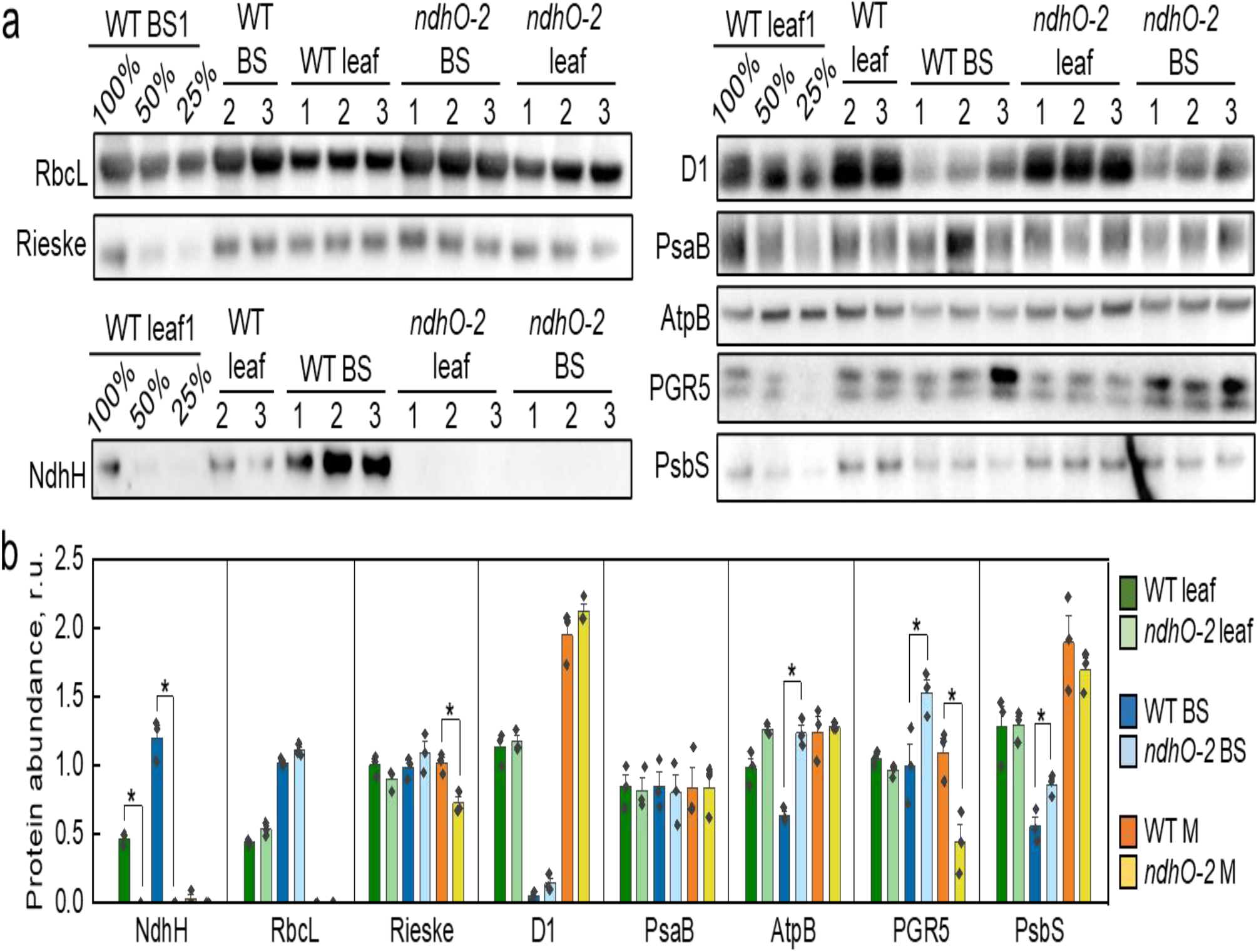
Abundance of photosynthetic proteins in wild type (WT) *S. viridis* and *ndhO-2* plants lacking NDH. **a**. Immunodetection of RbcL (the large subunit of Rubisco), Rieske (Cyt*b*_6_*f*), NdhH (NDH), D1 (PSII), PsaB (PSI), AtpB (ATP synthase), PGR5 (the lower band, see Fig S6 for details) and PsbS in bundle sheath (BS) or leaf protein samples loaded on Chl (*a* + *b*) basis. Three biological replicates (1, 2, 3) were loaded for each group and the titration series of one of the samples was used for relative quantification. **b**. Relative quantification of protein abundance per unit of Chl in leaves, BS and mesophyll (M) cells. Mean ± SE and three replicates are shown. Each protein has its own relative scale. Asterisks indicate statistically significant differences between the same cell types of WT and *ndhO-2* plants (*t*-test, *P* < 0.05).

### Electron fluxes in bundle sheath cells

Effects of the lack of NDH on BS electron transport were examined by membrane inlet O_2_ mass spectrometry and spectroscopy using isolated BS strands supplied with NaHCO_3_ and triose phosphate to support CO_2_ assimilation^30^. The rate of gross O_2_ evolution by PSII in *ndh* BS cells was double that of the WT BS (*P* = 0.037, Fig. 4a). Next, electron flux through PSI was compared by monitoring the absorbance signal of P700^+^, a cation of the reaction centre of PSI. The maximum oxidisable P700, P_M_, was similar between *ndhO* and WT plants (*P* = 0.55) indicating a similar amount of active PSI (Fig. 4b). These measured PSII and PSI activities were in line with relative abundances of D1 and PsaB in BS cells (Fig. 3).

**Fig. 4.**
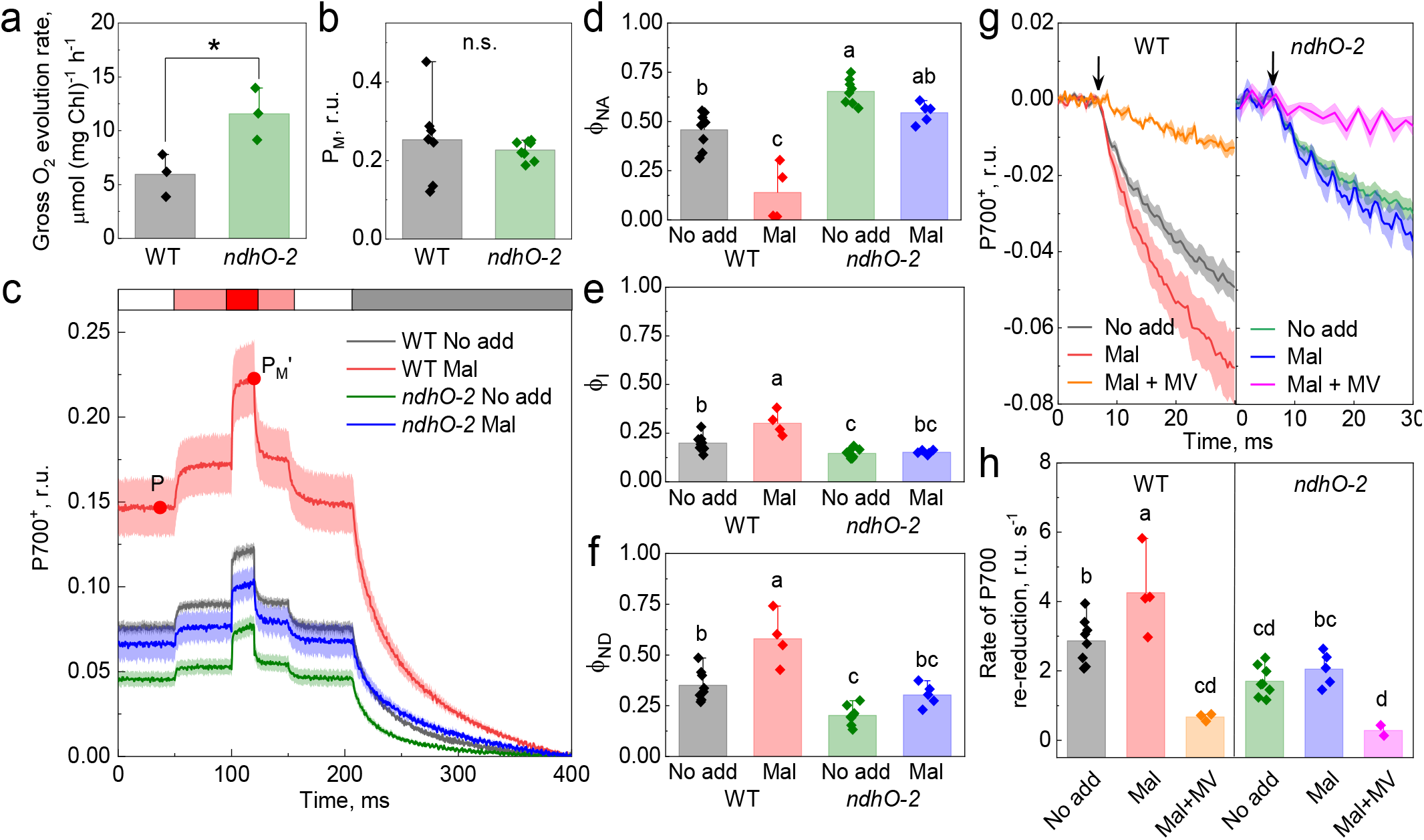
Photosynthetic properties of bundle sheath (BS) strands isolated from leaves of wild-type (WT) *S. viridis* and *ndhO-2* plants lacking NDH complex. All BS preparations were normalised to 25 μg Chl mL^-1^ and supplemented with NaHCO_3_ and triose phosphate. **a**. Gross O_2_ evolution rates of BS strands illuminated with white actinic light of 1000 μmol m^-2^ s^-1^ (*t*-test at *P* < 0.05). **b**. The maximum photo-oxidisable P700 (PSI reaction centre), P_M_ (n.s., not significant; *t*-test at *P* < 0.05). **c**. Fast kinetics of P700^+^ signal recorded from BS strands with or without 10 mM malate. BS were illuminated with white actinic light of 1000 μmol m^-2^ s^-1^ (white bar on top of the graph), actinic light with added far-red light (pink bar) or combined actinic, far-red light and a saturating pulse (red bar). Points on the WT trace with malate demonstrate how the P (the steady-state P700 oxidation level) and P_M_’ (the maximum level of P700 oxidation under light) values were obtained. **d**. The non-photochemical yield of PSI due to the acceptor side limitation (ϕ_NA_) calculated as (P_M_-P_M_’)/P_M_. **e**. The effective quantum yield of PSI (ϕ_I_) calculated as (P_M_’-P)/P_M_. **f**. The non-photochemical yield of PSI due to the donor side limitation (ϕ_ND_) calculated as P/P_M_. **g**. Fast kinetics of P700^+^ signal upon the light-dark shift normalised for the steady-state P700^+^ level for comparison of the kinetics. Arrows indicate the end of illumination. 200 μM of methyl viologen (MV) was added in addition to 10 mM of malate when indicated. **f**. The relative rate of P700 re-reduction estimated from initial slopes of the dark relaxation kinetics shown in (g). All bar graphs show Mean ± SD. (**c, g**) Traces are average of 2-5 biological replicates. Mean ± SE. (**d, e, f, h**) Letters indicate significant differences between the groups (one-way ANOVA and Tukey’s *post-hoc* test at α > 0.05).

The P700^+^ signal was next monitored from BS cells adapted to actinic light of 1000 μmol m^-2^ s^-1^ upon the application of strong far-red light and a saturating pulse allowing estimation of the photochemical yield of PSI (ϕ_I_) and the non-photochemical yields of the acceptor (ϕ_NA_) and donor (ϕ_ND_) sides of PSI (Fig. 4c). The traces were normalised at the minimal level of P700^+^ after a dark relaxation, and the steady-state (P) and maximum (P_M_’) levels of P700 oxidation under light were determined as shown in Fig. 4c. The addition of malate drastically reduced ϕ_NA_, *i*.*e*., the loss of PSI activity due to a lack of acceptors, in WT BS cells (*P* = 0.00003), compared to the value without malate (Fig. 4d). Concurrently, the addition of malate increased ϕ_I_ (by about 50%, *P* = 0.001) and ϕ_ND_ (*i*.*e*., the loss of PSI activity due to a lack of electrons on the donor side, *P* = 0.0003) in WT BS cells, compared to values without malate (Fig. 4e,f). BS cells of *ndh* plants showed 36% lower ϕ_I_ (*P* = 0.041) as well as higher ϕ_NA_ (*P* = 0.001) and lower ϕ_ND_ (*P* = 0.003) compared to WT BS cells already in the absence of malate. The addition of malate did not result in any significant changes in PSI activity in *ndh* BS cells (*P* = 0.99 for ϕ_I_, 0.16 for ϕ_NA_ and 0.12 for ϕ_ND_). Consequently, in the presence of malate, ϕ_I_ and ϕ_ND_ were about 50% lower than in WT BS cells (*P* = 0.00003 and *P* = 0.0001 respectively) while ϕ_NA_ was 4-fold higher (*P* = 0.000004) in *ndh* BS cells compared to WT.

The initial decay of P700^+^ signal upon the termination of actinic light provides an estimate of a relative rate of P700 re-reduction from the intersystem electron transport chain (Fig. 4g,h). Despite a WT-like Cyt*b*_6_*f* content (Fig. 3), *ndh* showed about 40% slower PSI reduction than WT plants in the absence of malate (*P* = 0.0128). The addition of malate accelerated the reduction of PSI in WT BS cells (*P* = 0.0158) but not in *ndh* BS cells (*P* = 0.92), compared to the rate without malate, resulting in about 50% lower reduction rate in the mutant compared to WT (*P* = 0.0003). The addition of methyl viologen (MV, a strong electron acceptor competing for electrons from PSI with CEF) together with malate allowed estimating the rate of P700 re-reduction by LEF only (Fig. 4g,h). In the presence of MV, the PSI reduction rate in BS cells was drastically reduced to about 16% and 13% in WT and *ndh*, respectively, compared to the rates observed with malate only.

### Leaf photosynthesis

Absence of NDH had significant impact on leaf electron transport and gas-exchange properties. Although the maximum photochemical efficiency of PSII (F_V_/F_M_) did not differ between the genotypes (Table S1), in *ndh* plants up to 50% less light absorbed by PSII was used for photochemical reactions (ϕ_II_) at all irradiances (Fig. 5a). The yield of regulated non-photochemical reactions (ϕ_NPQ_) was increased up to 2-fold in *ndh* leaves below 1600 μmol m^-2^ s^-1^, whilst the non-regulated non-photochemical yield (ϕ_NO_) was mostly unaltered, compared to WT (Fig. 5b and Fig. 5c). Due to the low PSII content in BS cells (Fig. 3), leaf chlorophyll a fluorescence originates mostly from mesophyll cells, whereas leaf P700^+^ absorbance reports on PSI activity in both mesophyll and BS cells. Plants lacking NDH had on average lower ϕ_I_ which was significantly reduced to about 50% of WT level at irradiances above 1500 μmol m^-2^ s^-1^ (Fig. 5d). The non-photochemical yields of PSI showed contrasting responses depending on irradiance. Below 200 μmol m^-2^ s^-1^, the loss of PSI photochemical efficiency in *ndh* was due to a lack of electron supply on the donor side whilst above 400 μmol m^-2^ s^-1^ it was due to the lack of acceptors (Fig. 5e and Fig. 5f).

**Fig. 5.**
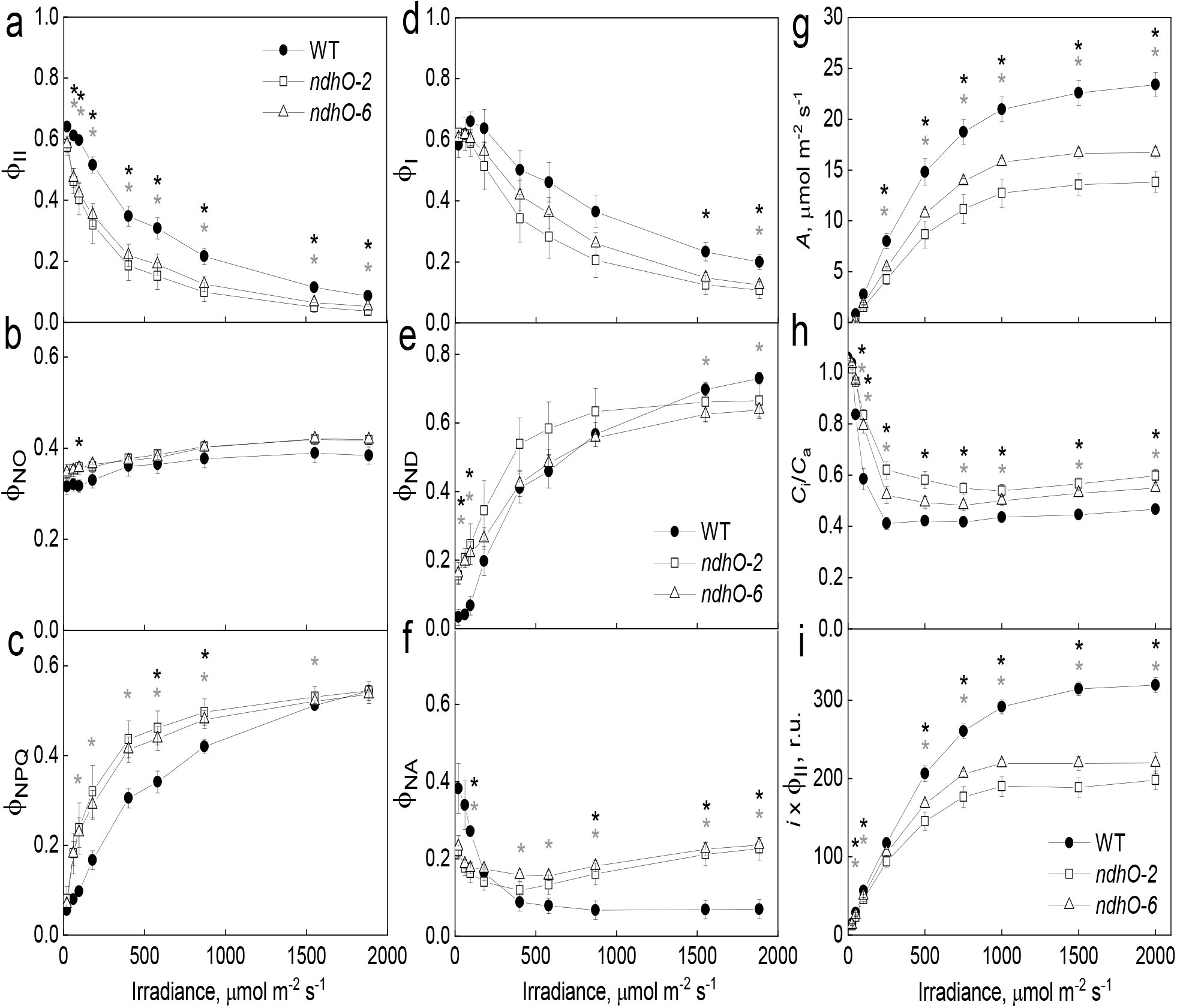
Leaf photosynthesis in wild-type *S. viridis* (WT) and gene-edited plants lacking NDH (*ndhO-2* and *ndhO-6*). (**a, b, c**) The photochemical yield of PSII (ϕ_II_), the yield of non-regulated energy dissipation (ϕ_NO_) and the yield of non-photochemical quenching (ϕ_NPQ_) in PSII. (**d, e, f**) The photochemical yield of PSI (ϕ_I_) and the non-photochemical yields of PSI donor (ϕ_ND_) and acceptor (ϕ_NA_) sides. PSI and PSII quantum yields were analysed concomitantly at different irradiance with Dual-PAM-100. (**g, h, i**) CO_2_ assimilation rate (*A*), the ratio of intercellular to ambient CO_2_ partial pressures (*C*_i_/*C*_a_) and a relative electron flux through PSII (*i* x ϕ_II_) measured concomitantly at different irradiance using Licor-6800. Mean ± SE, *n* = 4 biological replicates. Asterisks indicate statistically significant differences between the edited and WT plants (one-way ANOVA and Dunnett’s *post-hoc* test, α = 0.05): black asterisks for *ndhO-2*, grey asterisks for *ndhO-6*.

Plants lacking NDH exhibited about 50% lower CO_2_ assimilation rates at all irradiances (Fig. 5g) and lower quantum yield of CO_2_ assimilation, compared to WT (Table S1). The ratio of intercellular to ambient CO_2_ partial pressures (*C*_i_/*C*_a_), was significantly higher in *ndh* plants at all irradiances (Fig. 5h), compared to WT, indicating a higher CO_2_ availability at the site of primary carboxylation (*i*.*e*., in the cytosol of mesophyll cells). Calculated relative electron flux through PSII was significantly lower at all irradiances (Fig. 5i) whilst the slope of the relationship between NPQ and electron flux was drastically increased (Fig. S3) in plants lacking NDH in comparison to WT.

### Proton fluxes in mesophyll cells

Since no changes in electrochromic shift signal could be detected from the dark-adapted isolated BS cells in response to a saturating flash (Fig. S4a), leaf electrochromic shift signal was reported on thylakoid membrane energisation in mesophyll cells. Total *pmf*, which represents a balance between a build-up and dissipation of a transmembrane proton gradient, was similar between *ndh* and WT plants below 300 μmol m^-2^ s^-1^ and significantly decreased in the gene-edited plants at higher irradiances (Fig. 6a). Proton conductivity of the thylakoid membrane (*g*_H+_), indicative of the flux through ATP synthase, was significantly decreased in *ndh* leaves at 50 μmol m^-2^ s^-1^ but similar to WT at irradiances above that (Fig. 6b). The relationship between the light-driven proton flux through the thylakoid membrane (*ν*_H+_) and the relative electron flux through PSII (*i* x ϕ_II_), indicative of the CEF/LEF ratio^31^, remained constant between *ndh* and WT plants (Fig. 6c). However, the relationship between NPQ and *pmf* was markedly altered so that in *ndh* plants, for a given *pmf*, NPQ was approximately 2-fold higher than in WT (Fig. 6d). This altered relationship was not caused by changes in partitioning between ΔpH and Δψ (Fig. S4c and Fig. S4d). Moreover, a build-up of NPQ occurred at a similar rate upon illumination and, at 1 min after the termination of illumination, NPQ relaxed to a similar level in *ndh* and WT plants (Fig. S5).

**Fig. 6.**
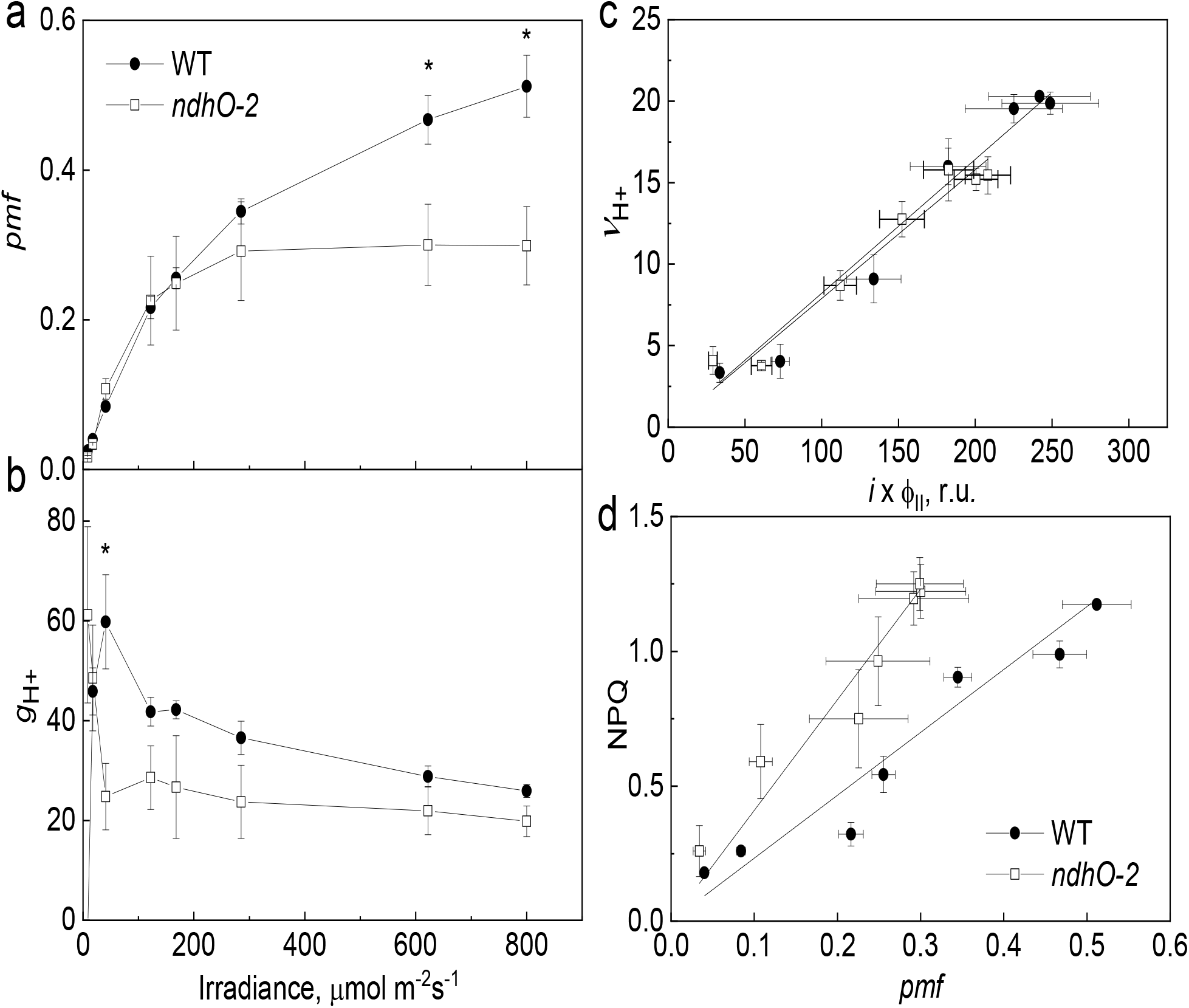
Thylakoid membrane energisation in wild-type (WT) *S. viridis* and gene-edited plants lacking NDH (*ndhO-2*). (**a, b**) Proton motive force (*pmf*) and proton conductivity of the thylakoid membrane (*g*_H+_) after 3-min illumination at different irradiance. **c**. The relationship between the light-driven proton flux (ν_H+_) and the relative electron flux through PSII. The slope of linear regression of ν_H+_ versus *i* x ϕ_II_ is 0.088 ±0.017 for WT and 0.080 ±0.012 for *ndhO-2* (*P* = 0.71). **d**. The relationship between the energy-dependent non-photochemical quenching (NPQ) and *pmf*. The slope of linear regression of NPQ versus *pmf* is 2.25 ±0.6 for WT and 4.14 ±0.33 for *ndhO-2* (*P* = 0.03). Mean ± SE, *n* = 4 biological replicates. Asterisks indicate statistically significant differences between the edited and WT plants at *P* < 0.05. The relationship between mean values of edited and WT plants was tested by the Student’s *t*-test.

## Discussion

NDH is believed to have evolved from the cyanobacterial NDH-1 complex^32^ and is largely conserved in the green lineages^33^. Despite being non-essential in normal conditions, due to its ability to maximise *pmf* during CEF^34^, NDH abundance increases in C_3_ plants in response to elevated ATP demand^35^ to facilitate plant acclimation to low or fluctuating light^36,37^. During the evolutionary transition from C_3_ to C_4_ plants, NDH levels increased concurrently with the emergence of C_4_-like species coinciding with the appearance of the C_4_ cycle as a CO_2_ pump^38^. Supporting an essential role of NDH in C_4_ photosynthesis, generated here *S. viridis* plants lacking NDH could not efficiently use absorbed light and reached only low assimilation rates (Fig. 2e and Fig. 5g). However, in contrast to mutants and transgenic plants with an impaired carbon concentrating mechanism^28,29^, growth of *ndh* plants did not recover at high CO_2_ (Fig. 2d). Therefore, NDH is not directly involved in the C_4_ cycle or concentrating CO_2_ in BS cells as previously proposed^25^; instead, our results indicate that NDH is essential for supporting CO_2_ assimilation by the C_3_ cycle.

Active CEF in BS cells has long been proposed to increase the ATP yield of C_4_ plants and contribute to fulfilling the energy demand of the carbon concentrating mechanism^15,39^. The measured CEF to LEF ratio (CEF/LEF) of C_4_ leaves of about 1.7 greatly exceeded the C_3_ leaf value of 0.9^40^ but no experimental estimates of CEF in BS cells were available. To study BS electron transport properties, here we isolated BS strands from leaves of *S. viridis* and supplied all preparations with triose phosphate and NaHCO_3_ to satisfy NADPH and (partially) CO_2_ requirements of the C_3_ cycle^30^. Thus, the electron transport chain of BS cells only needed to produce ATP to support C_3_ cycle activity. A strong ϕ_NA_ in WT BS cells in the absence of malate and its drastic decrease (combined with an increase of ϕ_ND_) after the malate addition corroborated that, without malate, activity of the C_3_ cycle was limited by CO_2_ availability and potentially also by NADPH. CO_2_ and NADPH derived from malate increased the demand of the C_3_ cycle for ATP which doubled PSI activity in WT BS cells (Fig. 4e). In those conditions, 86% of electrons reducing PSI belonged to CEF (Fig. 4h), producing a CEF/LEF ratio of >6 which closely resembled the model predictions^41^. These results provided long forthcoming experimental evidence for PSI driving active CEF in C_4_ BS cells set to supply ATP for the C_3_ cycle.

Detailed electron transport analysis of BS cells from *ndh* plants showed that NDH mediated a large portion of CEF and was the primary electron acceptor from PSI in BS cells. These conclusions were supported by a decreased PSI reduction capacity (Fig. 4h) and a strong PSI acceptor side limitation observed in BS cells of *ndh* plants (Fig. 4d). A 50% lower PSI activity in BS cells in the absence of NDH was proportional to the reduction of leaf CO_2_ assimilation rate (Fig. 2e and Fig. 5g) demonstrating the indispensable role of NDH-CEF in BS ATP production. Interestingly, even in the absence of NDH, most of the electrons reducing PSI belonged to CEF (Fig. 4h). This residual CEF could be linked to PGR5, as suggested by the increased PGR5 content in BS cells of *ndh* plants (Fig. 3a), and/or to other unknown CEF pathways. Molecular mechanisms of PGR5’s action are, however, still under debate^42^, and further investigation is required to determine whether it contributes to ATP production in BS cells or is mainly involved in photoprotection^27,43^. It is also worth mentioning that electron transport limitations detected in BS cells of *ndh* plants were unlikely to be caused by photosynthetic control. Since BS cells have low LEF (Fig. 4a,h), the C_3_ cycle is not the primary PSI electron acceptor and does not regulate PSI activity through NADPH consumption. Therefore, PSI (and hence CEF) activity in BS cells is likely regulated via photosynthetic control that restricts electron transport at Cyt*b*_6_*f* to match the capacity of CO_2_ assimilation reactions. In BS cells of *ndh* plants, electron transport was limited even in the presence of malate despite an apparent easing up of photosynthetic control observed in WT BS cells upon malate addition (Fig. 4) due to increased ATP demand upon providing CO_2_ and NADPH.

Alternatively, a lack of PSI stimulation in response to malate in BS cells of *ndh* plants could also be explained by inefficient malate decarboxylation *in vivo*. A supply of triose phosphate combined with an impaired ATP generation capacity of BS cells in the absence of NDH likely resulted in an elevated NADPH/NADP^+^ ratio and unavailability of NADP^+^ for malic enzyme. This could indicate a potential function of NDH in facilitating NADPH oxidation and using electrons derived from it for poising CEF. In C_3_ mesophyll chloroplasts, a respiration-like non-photochemical reduction of the PQ pool from stromal reductants by NDH is a part of the chlororespiratory pathway dissipating energy together with the plastidial terminal oxidase^44^. In C_4_ plants, electron donation from stromal donors to CEF is supported by observations of malate-driven carbon reduction in sorghum BS cells independent of PSII activity^45^ and by the capacity for reduction of PSI following malate addition to maize BS cells^46^. Since BS cells are effectively constantly light-limited due to shading by mesophyll^41,47^, driving CEF with minimal PSII engagement would maximise the light received by PSI potentially contributing to the observation that the highest quantum yields across all C_4_ decarboxylation types are seen in plants utilizing the NADP-ME photosynthetic mechanism^48^. However, the reverse reaction of the plant chloroplast ferredoxin:NADP^+^ oxidoreductase (FNR, Fig. 1), also required for reduction of the PQ pool from stromal NADPH, although theoretically possible and utilised *in vitro*^49^, is yet to be shown *in vivo*.

The mesophyll CEF/LEF ratio was not affected in *ndh* plants (Fig. 6c), which was comparable to the low impact of NDH deletion on CEF in C_3_ plants in optimal conditions^35^. Instead, lower mesophyll Chl content (Table S1), lower electron flux through PSII and higher NPQ (Fig. 5, Fig. S3) in *ndh* plants resembled effects observed in *Flaveria bidentis* with genetically reduced Rubisco abundance^50^. This observation suggested that the absence of NDH in the BS and inability of the C_3_ cycle to efficiently assimilate CO_2_ provided a negative feedback on the C_4_ cycle resulting in downregulation of mesophyll electron transport. Interestingly, since *pmf* in *ndh* plants was significantly lower than in WT at irradiances above 500 μmol m^−2^ s^−1^ (Fig. 6), there was an increased sensitivity of NPQ to *pmf* (Fig. 6d). An altered relationship between NPQ and ΔpH (or *pmf*) has been previously reported in some genetically modified plants, for instance, lacking or over-producing chloroplast NADPH-dependent thioredoxin reductase, or with increased Cyt*b*_6_*f* abundance^51,52^. These alterations are likely a result of perturbed stromal redox regulation and are mediated through the chloroplast thioredoxin system which directly targets multiple NPQ components^52,53^. The increased sensitivity of NPQ in *ndh* plants was therefore also in line with persistent mesophyll stroma overreduction due to accumulation of NADPH caused by a slow-down of the C_4_ cycle activity in response to C_3_ cycle limitation.

## Conclusion

In NADP-ME species, NADPH for the C_3_ cycle can be supplied from mesophyll cells through the influx of malate and triose phosphate whilst providing ATP for regeneration of ribulose 1,5-bisphosphate, the substrate of Rubisco, is required in the BS (Fig. 1). The BS electron transport chain therefore becomes specialised to primarily generate ATP by operating active CEF. Impaired CEF in the absence of NDH jeopardises the supply of ATP to the C_3_ cycle, thus, making NDH indispensable for C_4_ photosynthesis. Engineering a fully operational NADP-ME type C_4_ photosynthesis into C_3_ plants will require upregulating NDH abundance in BS cells to achieve the desired increases in assimilation and radiation use efficiency.

## Materials and Methods

### Construct assembly, transformation and selection of edited plants

*S. viridis* plants with null *ndhO* alleles were created using CRISPR/Cas9 gene-editing. The genomic and coding sequences of *S. viridis ndhO* (Sevir.5G467100) were obtained from Phytozome (https://phytozome-next.jgi.doe.gov). Two 19 nt gRNAs fitting the search criteria ‘A..19N..NGG’ within an exon, where NGG is the protospacer adjacent motif, were selected using CRISPOR^54^ (Fig. 2a). The use of ‘A’ for the first base of gRNAs was required to maximise the expression from *Oryza sativa* snoRNA *U3* (*OsU3*) promoter^55^. The *cas9* gene and *OsU3* promoter sequences were obtained from pRGEB32^56^ (Addgene) and adapted for the Golden Gate cloning system^57^. To create a gene construct for *ndhO* editing (Fig. 2b), the first expression module in a plant binary vector pAGM4723 was occupied by the hygromycin phosphotransferase gene (*hpt*) driven by the *O. sativa Actin1* (*OsAct1*) promoter. The second expression module contained *cas9* under the control of *Z. mays Ubiquitine1* (*ZmUbi1*) promoter. Both *hpt* and *cas9* were supplied with the bacterial nopaline synthase terminator. The third expression module in pAGM4723 was occupied by the two selected gRNAs forming a single synthetic polycistronic gene, assembled according to Xie, et al. ^56^, to be processed via the endogenous tRNA-processing system, under the control of *OsU3* promoter. The resulting construct was transformed into *S. viridis* cv. MEO V34-1 using the *Agrobacterium tumefaciens* strain AGL1^58^. T_0_ plants resistant to hygromycin were transferred to soil and analysed for *hpt* copy number by digital PCR (iDNA Genetics, Norwich, UK).

To select T_0_ plants with active Cas9, DNA was extracted from leaves with the DNeasy Plant kit (Qiagen, Venlo, The Netherlands) and the region of *ndhO* spanning both gRNAs was amplified using the primers CGCGTGGACAAGGAGAAGTA and CGTAGTCCAGCTTGTCCGAC. PCR products were sequenced and the edited *ndhO* alleles identified using Geneious Prime (https://www.geneious.com). Selected T_0_ plants were self-pollinated and their progeny analysed by digital PCR to retain only the plants that segregated out the T-DNA. Next, *ndhO* was sequenced to select the plants homozygous for *ndhO-2* and *ndhO-6* alleles (Fig. S1). The T_2_ progenies of those homozygous plants were used in all subsequent experiments. Gene and protein sequences were visualised in Geneious Prime, and protein structures were modelled using AlphaFold2^59^ and Mol*^60^.

### Plant growth conditions

All plants were grown in a controlled environment room with a 16 h light/8 h dark photoperiod, 28°C day, 22°C night, 60% humidity and ambient CO_2_ (if not stated otherwise) at a light intensity of 300 μmol m^−2^ s^−1^ supplied by 1000 W red sunrise 3200K lamps (Sunmaster Growlamps, Solon, OH). For the high CO_2_ growth experiment, air in the growth room was supplied with 2% CO_2_. For plants grown from seeds, seeds were surface-sterilised in 10% bleach and germinated on a rooting medium (pH 5.7) containing 2.15 g L^−1^ Murashige and Skoog salts, 10 ml L^−1^ 100× Murashige and Skoog vitamins stock, 30 g L^−1^ sucrose and 7 g L^−1^ Phytoblend. Seedlings that developed secondary roots were transferred to 0.6 L pots with garden soil mix layered on top with 2 cm of seed raising mix (Debco, Tyabb, Australia) both containing 1 g L^−1^ Osmocote (Scotts, Bella Vista, Australia). Wild-type (WT) *S. viridis* plants were used as control in all experiments. Youngest fully expanded leaves of the 3–4-week-old plants, before flowering, were used for all analyses.

### Chl analysis

Chl content in leaves, bundle sheath strands and protein samples was measured in 80% acetone buffered with 25 mM HEPES-KOH (pH 7.8) according to Porra, et al. ^61^. The portion of leaf Chl in BS cells (Chl_BS_ / Chl_leaf_) was calculated from the RbcL immunoblots as described in Ermakova, et al. ^14^. Chl contents of BS and mesophyll cells per leaf area (Chl_BS_ and Chl_MES_, Table S1) were calculated by multiplying leaf Chl content by the portion of leaf Chl in BS cells or in mesophyll.

### Blue-native PAGE, SDS-PAGE and immunoblotting

Thylakoid isolation and Blue-native PAGE were performed as described in Ermakova, et al. ^51^. For protein analysis, BS strands were isolated following the procedure of Ghannoum, et al. ^62^. Protein isolation from leaves and BS strands, SDS-PAGE and immunoblotting were performed as described in Ermakova, et al. ^14^. Aliquots were taken from protein samples for Chl analysis immediately after grinding. Samples were loaded either on leaf area or Chl (*a*+*b*) basis. Membranes were probed with antibodies against photosynthetic proteins: NdhH (1:3000, AS164065, Agrisera, Vännäs, Sweden), RbcL (1:10,000,^63^), PEPC (1:10,000,^64^), SBPase (1:3000, AS152873, Agrisera), D1 (1:10,000, AS10704, Agrisera), Rieske (1:5000, AS08330, Agrisera), PsaB (1:5000, AS10 695, Agrisera), AtpB (1:10,000, Agrisera), PGR5 (1:3000, AS163985, Agrisera, verification of this antibody is shown in Fig. S6), PsbS (1:3000, AS09533, Agrisera), Lhcb2 (1:10,000, AS01003, Agrisera). Quantification of immunoblots was performed with Image Lab software (Bio-Rad, Hercules, CA, USA). Relative protein abundances per unit of Chl in leaves (P_leaf_) and BS cells (P_BS_) were estimated from the immunoblots (Fig. 3). Using Chl contents of leaves and cells per leaf area (Chl_leaf_, Chl_BS_ and Chl_MES_ in Table S1), relative protein abundances per unit of Chl in mesophyll cells (P_MES_) were calculated as described in Ermakova, et al. ^14^ assuming that: P_leaf_ x Chl_leaf_ = P_BS_ x Chl_BS_ + P_MES_ x Chl_MES_.

### O_2_ evolution of BS strands

For activity assays, BS strands were isolated following the procedure of Furbank and Badger ^30^, resuspended in activity buffer (10 mM HEPES-KOH, pH 7.4, 2 mM MgCl_2_, 2 mM KH_2_PO_4_, 10 mM KCl, 0.3 M sorbitol) to a Chl (*a* + *b*) concentration of 25 μg mL^−1^, purged with N_2_ gas and kept on ice. Membrane inlet mass spectrometry (MIMS) was used to measure the steady-state gross O_2_ evolution rates through the production of ^16^O_2_ from suspensions of BS cells at a saturating irradiance (1000 μmol m^−2^ s^−1^). The system consisted of a magnetically stirred, thermally controlled at 25 °C sample cuvette separated from the high vacuum line of a Thermo Delta V ion ratio mass spectrometer (Thermo Electron Corp, Bremen, Germany) via a Teflon membrane (Hansatech Instruments, Norfolk, UK). The top lid of the cuvette had a quartz window through which halogen light was supplied through a fibre optic and a septum. BS suspensions were loaded into the cuvette, purged with N_2_ and then supplied with 1 mM HCO_3_ ^-^, 1 mg mL^-1^ carbonic anhydrase and 5 mM dihydroxyacetone phosphate to support CO_2_ assimilation^30^. Samples were left in darkness for 5 mins to reach equilibrium, followed by initiation of data acquisition for 2.5 min in the dark and 5 min at 1000 μmol m^-2^ s^-1^. Rates and calibrations were conducted as described in Beckmann, et al. ^65^.

### P700 spectroscopy on BS strands

For P700 measurements, 1 mL aliquot of BS suspension was incubated for five minutes at room temperature with 100 mM NaHCO_3_ and 5 mM dihydroxyacetone phosphate to support CO_2_ assimilation^30^ and with 10 mM malate and 200 μM methyl viologen when required. The suspension was then filtered via gentle vacuum onto a glass fiber disc (Whatman, Buckinghamshire, UK). The disc was saturated with activity buffer and metabolites and loaded into the measurement cuvette set at 25 °C. Redox changes of P700 were measured using the method and a set-up of Kou, et al. ^66^. A dual wavelength (820/870 nm) unit ED-P700DW (Heinz Walz, Effeltrich, Germany) was attached to a pulse amplitude modulation fluorometer PAM-101 (Heinz Walz) in the reflectance mode. The signal was zeroed in darkness before measurements were made. To determine P_M_, the maximal P700^+^ signal, samples without added malate were illuminated with weak far-red light of 30 μmol m^2^ s^-1^ to preferentially drive PSI photochemistry, over which a saturating pulse (10 μs at 9000 μmol m^2^ s^-1^) was triggered.

Next, samples were illuminated for three minutes with white actinic light of 1000 μmol m^2^ s^-1^ to reach a steady-state. The same actinic light was then maintained for 9.016 s, using an electronic shutter controlled by one terminal of a pulse/delay generator (Model 555, Berkeley Nucleonics, San Rafael, CA, USA). During the 9.016-s illumination cycle, data acquisition (using software written by the late AB Hope) was started by a second terminal of the pulse/delay generator yielding P, a steady-state P700^+^ level. At 8.85 s, a strong far-red light (2000 μmol m^-2^ s ^-1^) from an LED (741 nm ± 13 nm, LED735–66–60, Roithner LaserTechnik, Vienna, Austria) was triggered on for 100 ms using a third terminal of the pulse/delay generator to help oxidise P700^50^. While the far-red light was on, at 8.90 s, a saturating light pulse (9000 μmol m^-2^ s ^-1^) was applied for 20 ms from a fourth terminal of the pulse/delay generator, yielding P_M_’, the maximal oxidised level of P700 under light. Finally, the white actinic light was turned off by the electronic shutter at 9.016 s whilst the data acquisition continued for 85 ms. Immediately after a completion of one cycle of illumination and data acquisition, another 9.016-s cycle restarted, and sixteen traces were averaged automatically to improve the signal-to-noise ratio. The photochemical yield of PSI, ϕ_I_, was calculated as (P_M_’-P)/P_M_; the non-photochemical yield of the PSI acceptor side, ϕ_NA_, was calculated as (P_M_-P_M_’)/P_M_; the non-photochemical yield of PSI due to the donor side limitation, ϕ_ND_, was calculated as P/P_M_. The rate of P700 re-reduction was obtained from the initial slope of P700^+^ decline in the dark.

### Leaf spectroscopic and fluorescence analyses

Leaf PSII and PSI yields were measured with the Dual PAM-100 (Heinz Walz) under the red actinic light (635 nm). PSII activity was assessed with the pulse amplitude modulated fluorescence method using 620 nm measuring light of 9 μmol m^-2^ s^-1^. The redox state of P700 was assessed by detecting absorbance changes of the cation at 830 nm with a dual wavelength unit (830/875 nm). Saturating pulses of red light (635 nm) at 12,000 μmol m^-2^ s^-1^ were used. Leaves were first dark adapted for 30 min and then the maximum (F_M_) and the minimum (F_0_) levels of fluorescence in the dark were recorded upon the application of a saturating pulse. The maximum quantum yield of PSII (F_V_/F_M_) was calculated from those values as (F_M_-F_0_)/ F_M_. Then, the maximal P700^+^ signal (P_M_) was recorded upon the application of a saturating pulse at the end of the far-red light (720 nm) illumination, and the minimal P700^+^ signal (P_0_) was recorded after the saturating pulse. After that, leaves were illuminated for 10 min with actinic light of 400 μmol m^-2^ s^-1^ followed by 2 min of darkness, during which saturating pulses were applied every 40 s to probe for a build-up and relaxation of NPQ. NPQ was calculated after Bilger and Björkman ^67^ as (F_M_-F_M_’)/F_M_’ where F_M_’ is the maximum level of fluorescence under light recorded upon the application of a saturating pulse.

PSI and PSII parameters were then tested at 2-min intervals of increasing irradiance (0 to 1890 μmol m^-2^ s^-1^) by applying a saturating pulse at the end of each period. Recording F_M_’ and F (the fluorescence level briefly before the application of a saturating pulse) allowed tracking of the partitioning of absorbed excitation energy within PSII between the photochemical (ϕ_II_) and non-photochemical reactions, including the regulated (ϕ_NPQ_) and non-regulated (ϕ_NO_) fractions^68^. Using the steady-state P700^+^ signal (P) and the maximal P700^+^ signal under light (P_M_′) recorded just before and upon the pulse application, respectively, we calculated the photochemical yield of PSI (ϕ_I_) and the non-photochemical yields of PSI due the acceptor (ϕ_NA_) or donor (ϕ_ND_) side limitations^69^.

### Gas-exchange analysis

Concomitant gas exchange and fluorescence analyses were performed at different irradiances and intercellular CO_2_ partial pressures using a LI-6800 (LI-COR Biosciences, Lincoln, NE, USA) equipped with a fluorometer head 6800-01A (LI-COR Biosciences). 90% red/10% blue actinic light was used for all measurements. First, leaves were equilibrated at 381 μbar CO_2_ in the reference side, leaf temperature 25°C, 60% humidity, and a flow rate of 500 μmol s^−1^. Then, either CO_2_ partial pressures from 0 to 1200 μbar were imposed at 3-min intervals at a constant irradiance of 1500 μmol m^−2^ s^−1^ (CO_2_ response curve) or a stepwise increase of irradiance from 0 to 2000 μmol m^−2^ s^−1^ at 2-min intervals was imposed at a constant 381 μbar of CO_2_ in the reference side (light response curve). A relative electron flux through PSII (*i* x ϕ_II_) was estimated upon the application of multiphase saturating pulses (8000 μmol m^−2^ s^−1^) as irradiance multiplied by the photochemical yield of PSII^70^.

### Thylakoid membrane energisation

Electrochromic shift signal (ECS) was monitored as absorbance changes between 550 and 515 nm with the Dual PAM-100 equipped with the P515/535 emitter-detector module (Heinz Walz) and normalised for the amplitude of ECS response to a saturating pulse (20 ms, 14,000 μmol m^−2^ s^−1^) measured from the dark-adapted leaves. Measurements were conducted on dark-adapted for 40 min leaves during 3-min light and 3-min dark intervals of increasing irradiance. Proton motive force (*pmf*) and ΔpH were estimated upon the shift from light to dark (Fig. S4)^71^. Proton conductivity of the thylakoid membrane (*g*_H+_) was calculated as an inverse time constant of the first order exponential ECS decay^70^ fitted in OriginPro 2018b (OriginLab, Northampton, MA, USA), and the light-driven proton flux (*ν*_H+_) was calculated as the initial rate of change in the ECS signal upon light termination^72^.

### Statistical analysis

ANOVA and two-tailed heteroscedastic Student’s *t*-test were performed in OriginPro 2018b. Details of replication, *post-hoc* tests and *P* values are provided in the main text and in figure legends.

## Acknowledgements

We thank Brendon Conlan and Spencer Whitney for a gift of vectors and antibodies, Fred Chow for equipment and assistance with P700 measurements, Xueqin Wang for Setaria transformation, and Emily Watson, Samuel James Nix and Zac Taylor for technical assistance. This work was supported by the Australian Research Council Centre of Excellence for Translational Photosynthesis (CE140100015).

## References

1 Sage, R. F. & Zhu, X.-G. Exploiting the engine of C_4_ photosynthesis. Journal of Experimental Botany 62, 2989–3000 (2011).

2 Long, S. P. Environmental responses. C_4_ plant biology, 215-249 (1999).

3 Hatch, M. D. C_4_ photosynthesis - a unique blend of modified biochemistry, anatomy and ultrastructure. Biochimica Et Biophysica Acta 895, 81–106 (1987).

4 Furbank, R. T. Evolution of the C_4_ photosynthetic mechanism: are there really three C_4_ acid decarboxylation types? Journal of Experimental Botany 62, 3103–3108 (2011). 10.1093/jxb/err080

5 Ermakova, M. et al. Installation of C_4_ photosynthetic pathway enzymes in rice using a single construct. Plant Biotechnology Journal 19, 575–588 (2021).

6 Ermakova, M., Danila, F. R., Furbank, R. T. & von Caemmerer, S. On the road to C_4_ rice: advances and perspectives. The Plant Journal 101, 940–950 (2020).

7 Munekage, Y. Light harvesting and chloroplast electron transport in NADP-malic enzyme type C_4_ plants. Current Opinion in Plant Biology 31, 9–15 (2016).

8 Malone, L. A., Proctor, M. S., Hitchcock, A., Hunter, C. N. & Johnson, M. P. Cytochrome b6f – orchestrator of photosynthetic electron transfer. Biochimica et Biophysica Acta (BBA) - Bioenergetics 1862, 148380 (2021).10.1016/j.bbabio.2021.148380

9 Müller, P., Li, X.-P. & Niyogi, K. K. Non-photochemical quenching. A response to excess light energy. Plant physiology 125, 1558–1566 (2001).

10 Li, X.-P. et al. Regulation of photosynthetic light harvesting involves intrathylakoid lumen pH sensing by the PsbS protein. Journal of Biological Chemistry 279, 22866–22874 (2004).

11 Demmig-Adams, B. Carotenoids and photoprotection in plants: A role for the xanthophyll zeaxanthin. Biochimica et Biophysica Acta (BBA) - Bioenergetics 1020, 1–24 (1990).

12 Kramer, D. M. & Evans, J. R. The importance of energy balance in improving photosynthetic productivity. Plant Physiology 155, 70–78 (2011).

13 von Caemmerer, S. & Furbank, R. T. Strategies for improving C_4_ photosynthesis. Current Opinion in Plant Biology 31, 125–134 (2016).

14 Ermakova, M. et al. Upregulation of bundle sheath electron transport capacity under limiting light in C_4_ Setaria viridis. The Plant Journal 106, 1443–1454 (2021).

15 Furbank, R., Jenkins, C. & Hatch, M. C_4_ photosynthesis: quantum requirement, C_4_ and overcycling and Q-cycle involvement. Functional Plant Biology 17, 1–7 (1990).

16 Woo, K. C., Gerbaud, A. & Furbank, R. T. Evidence for endogenous cyclic photophosphorylation in intact chloroplasts: ^14^CO_2_ fixation with dihydroxyacetone phosphate. Plant Physiology 72, 321–325 (1983).

17 Shikanai, T. Cyclic Electron Transport Around Photosystem I: Genetic Approaches. Annual Review of Plant Biology 58, 199–217 (2007). 10.1146/annurev.arplant.58.091406.110525

18 Munekage, Y. et al. Elevated expression of PGR5 and NDH-H in bundle sheath chloroplasts in C_4_ Flaveria species. Plant and Cell Physiology 51, 664–668 (2010).

19 Munekage, Y. & Taniguchi, Y. Y. Promotion of cyclic electron transport around Photosystem I with the development of C_4_ photosynthesis. Plant and Cell Physiology 57, 897–903 (2016).

20 Majeran, W. et al. Consequences of C_4_ differentiation for chloroplast membrane proteomes in maize mesophyll and bundle sheath cells. Molecular & Cellular Proteomics 7, 1609–1638 (2008).

21 Kubicki, A., Funk, E., Westhoff, P. & Steinmüller, K. Differential expression of plastome-encoded ndh genes in mesophyll and bundle-sheath chloroplasts of the C_4_ plant Sorghum bicolor indicates that the complex I-homologous NAD(P)H-plastoquinone oxidoreductase is involved in cyclic electron transport. Planta 199, 276–281 (1996).

22 Takabayashi, A., Kishine, M., Asada, K., Endo, T. & Sato, F. Differential use of two cyclic electron flows around photosystem I for driving CO_2_-concentration mechanism in C_4_ photosynthesis. Proceedings of the National Academy of Sciences of the United States of America 102, 16898–16903 (2005).

23 Sazanov, L. A., Burrows, P. & Nixon, P. J. Detection and characterization of a complex I-like NADH-specific dehydrogenase from pea thylakoids. Biochemical Society Transactions 24, 739–743 (1996).

24 Shikanai, T. et al. Directed disruption of the tobacco ndhB gene impairs cyclic electron flow around photosystem I. Proceedings of the National Academy of Sciences 95, 9705–9709 (1998).

25 Peterson, R. B., Schultes, N. P., McHale, N. A. & Zelitch, I. Evidence for a role for NAD(P)H dehydrogenase in concentration of CO_2_ in the bundle sheath cell of Zea mays. Plant Physiology 171, 125 (2016).

26 Ishikawa, N. et al. NDH-mediated cyclic electron flow around Photosystem I is crucial for C_4_ photosynthesis. Plant and Cell Physiology 57, 2020–2028 (2016).

27 Ogawa, T. et al. Two cyclic electron flows around photosystem I differentially participate in C_4_ photosynthesis. Plant Physiology (2023). 10.1093/plphys/kiad032

28 Alonso-Cantabrana, H. et al. Diffusion of CO_2_ across the mesophyll-bundle sheath cell interface in a C_4_ plant with genetically reduced PEP carboxylase activity. Plant Physiology 178, 72–81 (2018).

29 Danila, F. R. et al. Bundle sheath suberisation is required for C_4_ photosynthesis in a Setaria viridis mutant. Communications Biology 4, 254 (2021).

30 Furbank, R. T. & Badger, M. R. Photorespiratory characteristics of isolated bundle sheath strands of C_4_ monocotyledons. Functional Plant Biology 10, 451–458 (1983).

31 Baker, N. R., Harbinson, J. & Kramer, D. M. Determining the limitations and regulation of photosynthetic energy transduction in leaves. Plant, Cell & Environment 30, 1107–1125 (2007). 10.1111/j.1365-3040.2007.01680.x

32 Laughlin, T. G., Savage, D. F. & Davies, K. M. Recent advances on the structure and function of NDH-1: The complex I of oxygenic photosynthesis. Biochimica et Biophysica Acta (BBA) - Bioenergetics 1861, 148254 (2020).

33 Ruhlman, T. A. et al. NDH expression marks major transitions in plant evolution and reveals coordinate intracellular gene loss. BMC Plant Biology 15, 100 (2015).

34 Strand, D. D., Fisher, N. & Kramer, D. M. The higher plant plastid NAD(P)H dehydrogenase-like complex (NDH) is a high efficiency proton pump that increases ATP production by cyclic electron flow. Journal of Biological Chemistry 292, 11850–11860 (2017).

35 Livingston, A. K., Cruz, J. A., Kohzuma, K., Dhingra, A. & Kramer, D. M. An Arabidopsis mutant with high cyclic electron flow around Photosystem I (hcef) Involving the NADPH dehydrogenase complex The Plant Cell 22, 221–233 (2010).

36 Yamori, W., Makino, A. & Shikanai, T. A physiological role of cyclic electron transport around photosystem I in sustaining photosynthesis under fluctuating light in rice. Scientific Reports 6 (2016). 10.1038/srep20147

37 Nakano, H., Yamamoto, H. & Shikanai, T. Contribution of NDH-dependent cyclic electron transport around photosystem I to the generation of proton motive force in the weak mutant allele of pgr5. Biochimica et Biophysica Acta (BBA) - Bioenergetics 1860, 369–374 (2019). 10.1016/j.bbabio.2019.03.003

38 Lyu, M.-J. A. et al. The coordination of major events in C_4_ photosynthesis evolution in the genus Flaveria. Scientific Reports 11, 15618 (2021).

39 von Caemmerer, S. Updating the steady state model of C_4_ photosynthesis. Journal of Experimental Botany 72, 6003–6017 (2021). 10.1101/2021.03.13.435281

40 Sagun, J. V., Badger, M. R., Chow, W. S. & Ghannoum, O. Cyclic electron flow and light partitioning between the two photosystems in leaves of plants with different functional types. Photosynthesis Research 142, 321–334 (2019). 10.1007/s11120-019-00666-1

41 Bellasio, C. & Ermakova, M. Reduction of bundle sheath size boosts cyclic electron flow in C_4_ Setaria viridis acclimated to low light. The Plant Journal 111, 1223–1237 (2022).

42 Kanazawa, A. et al. Chloroplast ATP synthase modulation of the thylakoid proton motive force: implications for Photosystem I and Photosystem II photoprotection. Frontiers in Plant Science 8 (2017). 10.3389/fpls.2017.00719

43 Tazoe, Y. et al. Overproduction of PGR5 enhances the electron sink downstream of Photosystem I in a C_4_ plant, Flaveria bidentis. The Plant Journal 103, 814–823 (2020).

44 Peltier, G., Aro, E.-M. & Shikanai, T. NDH-1 and NDH-2 plastoquinone reductases in oxygenic photosynthesis. Annual Review of Plant Biology 67, 55–80 (2016). 10.1146/annurev-arplant-043014-114752

45 Osmond, B. Carbon reduction and photosystem II deficiency in leaves of C_4_ plants. Aust J Plant Physiol, 41-50 (1974).

46 Ivanov, B., Asada, K., Kramer, D. M. & Edwards, G. Characterization of photosynthetic electron transport in bundle sheath cells of maize. I. Ascorbate effectively stimulates cyclic electron flow around PSI. Planta 220, 572–581 (2005).

47 Bellasio, C. & Lundgren, M. R. Anatomical constraints to C_4_ evolution: light harvesting capacity in the bundle sheath. New Phytologist 212, 485–496 (2016).

48 Ehleringer, J. & Pearcy, R. W. Variation in Quantum Yield for CO2 Uptake among C3 and C4 Plants. Plant Physiology 73, 555–559 (1983). 10.1104/Pp.73.3.555

49 Munekage, Y. et al. PGR5 is involved in cyclic electron flow around Photosystem I and is essential for photoprotection in Arabidopsis. Cell 110, 361–371 (2002).

50 Siebke, K., von Caemmerer, S., Badger, M. & Furbank, R. T. Expressing an rbcS antisense gene in transgenic Flaveria bidentis leads to an increased quantum requirement for CO_2_ fixed in Photosystems I and II. Plant Physiology 115, 1163–1174 (1997).

51 Ermakova, M., Lopez-Calcagno, P. E., Raines, C. A., Furbank, R. T. & von Caemmerer, S. Overexpression of the Rieske FeS protein of the Cytochrome b_6_f complex increases C_4_ photosynthesis in Setaria viridis. Communications Biology 2 (2019).

52 Nikkanen, L., Guinea Diaz, M., Toivola, J., Tiwari, A. & Rintamäki, E. Multilevel regulation of non-photochemical quenching and state transitions by chloroplast NADPH-dependent thioredoxin reductase. Physiologia Plantarum 166, 211–225 (2019). 10.1111/ppl.12914

53 Da, Q. et al. M-type thioredoxins are involved in the xanthophyll cycle and proton motive force to alter NPQ under low-light conditions in Arabidopsis. Plant Cell Reports 37, 279–291 (2018). 10.1007/s00299-017-2229-6

54 Concordet, J.-P. & Haeussler, M. CRISPOR: intuitive guide selection for CRISPR/Cas9 genome editing experiments and screens. Nucleic Acids Research 46, 242–245 (2018).

55 Hassan, M. M. et al. Construct design for CRISPR/Cas-based genome editing in plants. Trends in Plant Science 26, 1133–1152 (2021).

56 Xie, K., Minkenberg, B. & Yang, Y. Boosting CRISPR/Cas9 multiplex editing capability with the endogenous tRNA-processing system. Proceedings of the National Academy of Sciences 112, 3570–3575 (2015).

57 Engler, C. et al. A Golden Gate modular cloning toolbox for plants. ACS Synthetic Biology 3, 839–843 (2014).

58 Osborn, H. L. et al. Effects of reduced carbonic anhydrase activity on CO_2_ assimilation rates in Setaria viridis: a transgenic analysis. Journal of Experimental Botany 68, 299–310 (2016).

59 Jumper, J. et al. Highly accurate protein structure prediction with AlphaFold. Nature 596, 583–589 (2021).

60 Sehnal, D. et al. Mol* Viewer: modern web app for 3D visualization and analysis of large biomolecular structures. Nucleic Acids Research 49, 431–437 (2021).

61 Porra, R. J., Thompson, W. A. & Kriedemann, P. E. Determination of accurate extinction coefficients and simultaneous equations for assaying chlorophylls a and b extracted with four different solvents: verification of the concentration of chlorophyll standards by atomic absorption spectroscopy. Biochimica et Biophysica Acta (BBA) - Bioenergetics 975, 384–394 (1989).

62 Ghannoum, O. et al. Faster rubisco is the key to superior nitrogen-use efficiency in NADP-malic enzyme relative to NAD-malic enzyme C_4_ grasses. Plant Physiology 137, 638–650 (2005).

63 Martin-Avila, E. et al. Modifying plant photosynthesis and growth via simultaneous chloroplast transformation of Rubisco large and small subunits. The Plant Cell 32, 2898–2916 (2020).

64 Karki, S. et al. A role for neutral variation in the evolution of C_4_ photosynthesis. bioRxiv (2020). 10.1101/2020.05.19.104299

65 Beckmann, K., Messinger, J., Badger, M. R., Wydrzynski, T. & Hillier, W. On-line mass spectrometry: membrane inlet sampling. Photosynthesis research 102, 511–522 (2009).

66 Kou, J. et al. Estimation of the steady-state cyclic electron flux around PSI in spinach leaf discs in white light, CO_2_-enriched air and other varied conditions. Functional Plant Biology 40, 1018–1028 (2013).

67 Bilger, W. & Björkman, O. Role of the xanthophyll cycle in photoprotection elucidated by measurements of light-induced absorbance changes, fluorescence and photosynthesis in leaves of Hedera canariensis. Photosynthesis Research 25, 173–185 (1990).

68 Kramer, D., Johnson, G., Kiirats, O. & Edwards, G. New fluorescence parameters for the determination of QA redox state and excitation energy fluxes. Photosynthesis Research 79, 209–218 (2004).

69 Klughammer, C. & Schreiber, U. Saturation pulse method for assessment of energy conversion in PS I. PAM Application notes 1, 3 (2008).

70 Sacksteder, C. A., Kanazawa, A., Jacoby, M. E. & Kramer, D. M. The proton to electron stoichiometry of steady-state photosynthesis in living plants: a proton-pumping Q cycle is continuously engaged. Proceedings of the National Academy of Sciences 97, 14283–14288 (2000).

71 Takizawa, K., Cruz, J. A., Kanazawa, A. & Kramer, D. M. The thylakoid proton motive force in vivo. Quantitative, non-invasive probes, energetics, and regulatory consequences of light-induced pmf. Biochimica et Biophysica Acta (BBA) - Bioenergetics 1767, 1233–1244 (2007).

72 Takizawa, K., Kanazawa, A. & Kramer, D. M. Depletion of stromal Pi induces high ‘energy-dependent’ antenna exciton quenching (qE) by decreasing proton conductivity at CFO-CF1 ATP synthase. Plant, Cell & Environment 31, 235–243 (2008).

