## Supplementary figures for "NDH complex-mediated cyclic electron flow in bundle sheath cells enables C_4_ photosynthesis"

### Supplementary materials

**Table S1**. Biochemical properties of wild type (WT) *S. viridis* and gene-edited plants lacking NDH. Mean ± SE, *n* = 3-4 biological replicates. BS, bundle sheath cells; M, mesophyll cells; N/A, not assessed; $F_V/F_M$, the maximum quantum yield of PSII. Asterisks indicate statistically significant differences between edited and WT plants (one-way ANOVA and Tukey's *post-hoc* test at α = 0.05 or Student's *t*-test at $P < 0.05$).

| Parameter | WT | *ndhO-2* | *ndhO-6* |
|---|---|---|---|
| $Chl_{leaf}$, mmol $m^{-2}$ | 0.48 ±0.03 | 0.34 ±0.05* | 0.36 ±0.05* |
| $Chl_{BS}$ / $Chl_{leaf}$ | 0.43 ±0.02 | 0.50 ±0.03 | N/A |
| $Chl_{BS}$, mmol $m^{-2}$ leaf | 0.21 ±0.03 | 0.17 ±0.01 | N/A |
| $Chl_M$, mmol $m^{-2}$ leaf | 0.27 ±0.03 | 0.17 ±0.03* | N/A |
| $F_V/F_M$ | 0.76 ±0.006 | 0.73 ±0.012 | 0.75 ±0.005 |
| Quantum yield, mol $CO_2$ (mol incident quanta)$^{-1}$ | 0.039 ±0.002 | 0.020 ±0.002* | 0.027 ±0.002* |

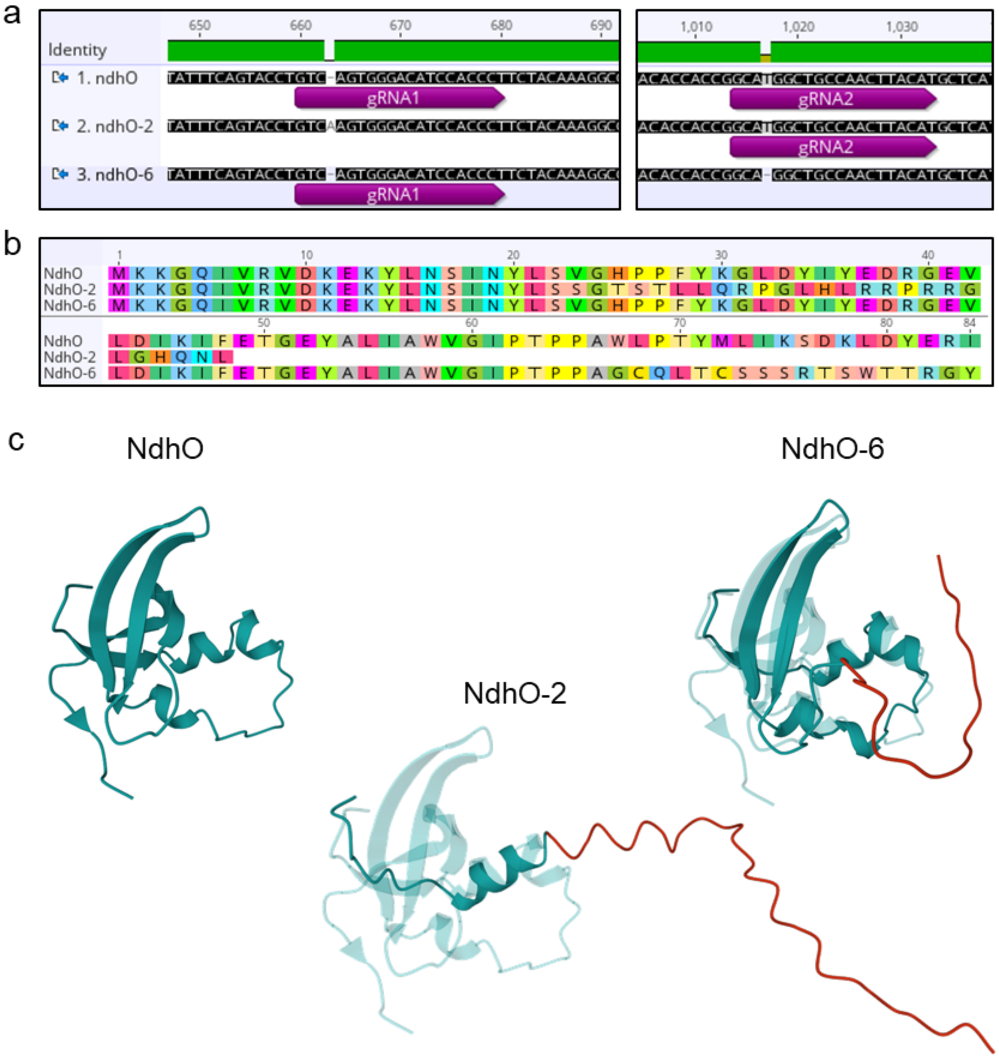


**Fig. S1**. New *ndhO* allelles created by CRISPR/Cas9 gene-editing in *S. viridis*. **a**. Sequencing of *ndhO* from a wild type (WT) plant and from the plants carrying homozygous edited alleles *ndhO-2* and *ndhO-6. ndhO-2* has a single nucleotide insertion at gRNA1 and *ndhO-6* has a single nucleotide deletion at gRNA2. **b**. Comparison of amino acid (aa) sequences encoded by *ndhO* (WT), *ndhO-2* and *ndhO-6* alleles. **c**. Protein structures of NdhO, NdhO-2 and NdhO-6 modelled based on aa sequence using AlphaFold2 and Mol*. NdhO-2 and NdhO-6 structures are aligned with and compared against NdhO (transparent teal), with aligned residues displayed in teal and misaligned residues displayed in red.

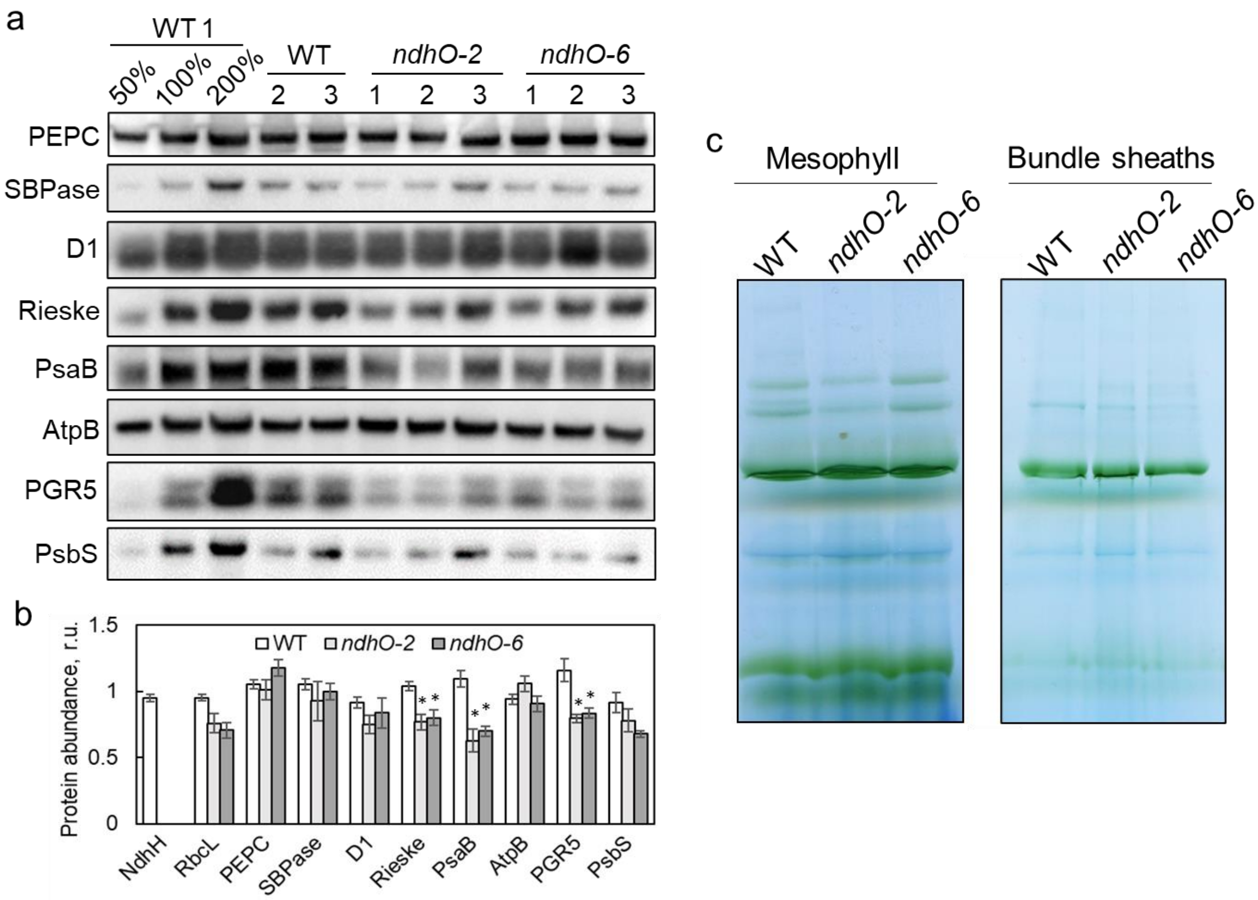


**Fig. S2**. Protein analysis of WT *S. viridis* and the gene-edited plants lacking NDH. **a**. Immunodetection of PEPC (PEP carboxylase), SBPase (sedoheptulose-bisphosphatase), D1 (PSII subunit), Rieske (Cyt$b_6f$ subunit), PsaB (PSI subunit), AtpB (ATP synthase subunit), PGR5 (the lower band, see Fig. S6) and PsbS in leaf protein extracts loaded on leaf area basis. Three biological replicates were loaded for each genotype and the titration series of one of the WT samples was used for relative quantification. **b**. Relative quantification of protein abundances from Fig. 2b and Fig. S2a per leaf area. Mean ± SE, $n$ = 3 biological replicates. Each protein has its own relative scale. Asterisks indicate statistically significant differences between edited and WT plants (one-way ANOVA and Dunnett's post-hoc test, $\alpha$ = 0.05). **c**. Thylakoid protein complexes separated by Blue-Native PAGE. 10 µg of Chl ($a$+$b$) loaded for each sample.

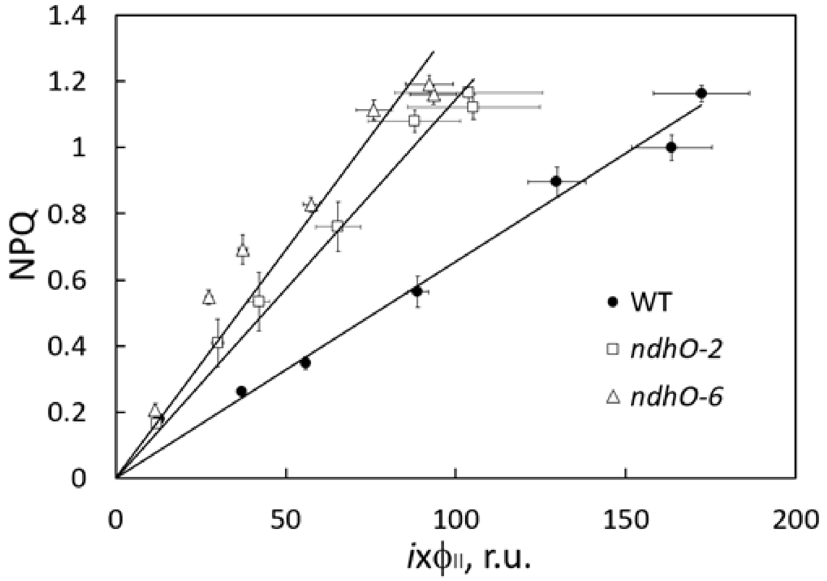


**Fig. S3**. A relationship between NPQ and the relative electron flux through PSII ($i$ x $\phi_{II}$) in WT *S. viridis* and gene-edited plants lacking NDH.

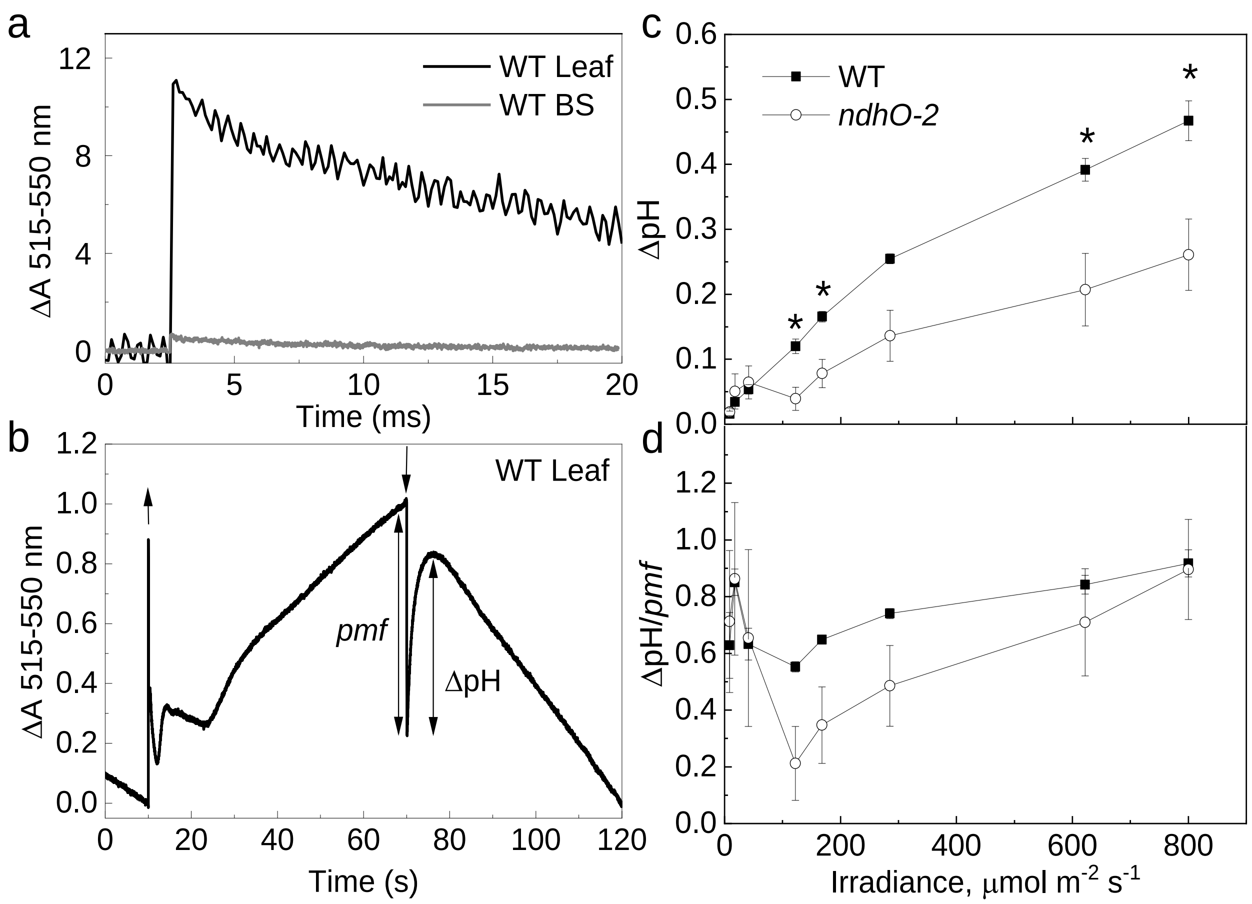


**Fig. S4**. Analysis of electrochromic shift signal (ECS) from leaves and bundle sheath (BS) cells of WT *S. viridis* and gene-edited plants lacking NDH (*ndhO-2*). **a.** Traces of saturating flash-induced ECS from leaves and isolated BS strands. BS sample of 25 µg Chl (a + b) was compared to leaf area containing about 4.2 µg Chl (a + b). **b.** Example of ECS trace recorded from WT leaf. Up and down arrows mark the beginning and the end of illumination with red actinic light of 500 µmol $m^{-2}$ $s^{-1}$. Double sided arrows indicate the amplitude of proton motive force (*pmf*) and ΔpH according to the parsing method[71]. (**c** and **d**) ΔpH and ΔpH/*pmf* at different irradiances. Asterisks indicate statistically significant differences between *ndhO-2* and WT (Student's t-test, $P < 0.05$).

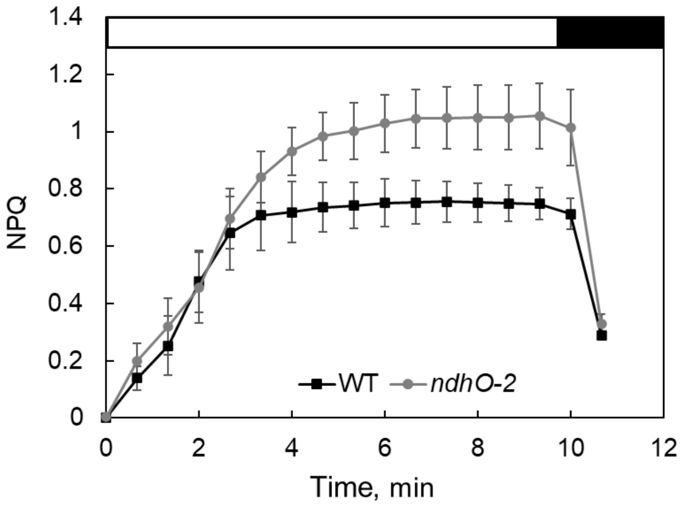


**Fig. S5**. A build-up and relaxation of NPQ in dark-adapted leaves of WT *S. viridis* and gene-edited plants lacking NDH. White bar, red actinic light at 400 µmol $m^{-2}$ $s^{-1}$; black bar, darkness.

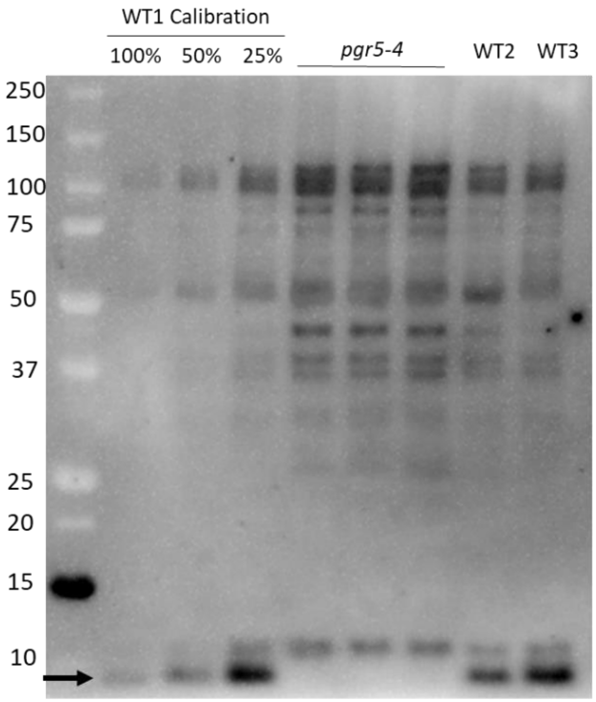


**Fig. S6**. Verification of the PGR5 antibody using protein extracts from leaves of wild-type (WT) *S. viridis* and gene-edited plants lacking PGR5. The dilution series of WT1 sample and three biological replicates for each group were loaded on leaf area basis. Arrow points to a band missing on the *pgr5* knock-out line which was used for quantification in Fig. 3.
